# Structural basis of nucleosomal H4K20 recognition and methylation by SUV420H1 methyltransferase

**DOI:** 10.1101/2023.07.27.549245

**Authors:** Folan Lin, Ruxin Zhang, Weihan Shao, Cong Lei, Ying Zhang, Mingxi Ma, Zengqi Wen, Wanqiu Li

## Abstract

Histone lysine methyltransferase of SUV420H1, which is responsible for site-specific di-/tri-methylation of histone H4 lysine 20 (H4K20), has crucial roles in DNA-templated processes, including DNA replication, DNA damage repair and chromatin compaction. Its mutations frequently occur in human cancers. Nucleosomes containing the histone variant H2A.Z enhance the catalytic activities of SUV420H1 in H4K20 di-methylation deposition, regulating early replication origins. However, the molecular mechanism of how H4K20 methyl marks are deposited on nucleosomes by SUV420H1 remains poorly understood. Here we report the cryo-electron microscopy (cryo-EM) structures of SUV420H1 associated with canonical nucleosome core particles (NCPs) or H2AZ containing NCPs. We find that SUV420H1 make extensive site-specific contacts with histone and DNA regions. SUV420H1 C-terminal domain recognizes the H2A-H2B acidic pocket of NCPs through its two arginine anchors, thus enabling H4K20 insertion for catalysis specifically. We also identify important residues increasing the catalytic activities of SUV420H1 bound to H2A.Z NCPs. In vitro and in vivo functional analysis reveal that multiple disease associated mutations at the interfaces are essential for its catalytic activity and chromatin states regulation. Together, our study provides molecular insights into the nucleosome-based recognition and methylation mechanisms of SUV420H1, and a structural basis for understanding SUV420H1-related human disease.

## Introduction

As the building block of chromatin, nucleosome is composed of 147bp DNA wraps around a compact histone octamer^1^, where the core histones possess a multiplicity of post-translational modifications (PTMs)^2^. Histone lysine methylation is a unique PTM because of its relative stability and multivalency, and plays a crucial role in a wide variety of DNA templated processes including transcription, replication and DNA repair^3, 4^. The canonical lysine methylation mark on histone H4 at lysine 20 (H4K20) is conserved from yeast to human. Histone H4 lysine 20 methylation (H4K20me) is a crucial modification for genome maintenance involved in replication, DNA damage repair and chromatin compaction^5–10^. There are three distinct H4K20 methylation states: mono-(H4K20me1), di-(H4K20me2) and trimethylation (H4K20me3), associated with different biological functions. It is known that SET8 (also known as PR-Set7) is responsible for the H4K20 mono-methylation (H4K20me1), whereas SUV420H1 and SUV420H2 catalyze H4K20 di-methylation (H4K20me2) and tri-methylation (H4K20me3). H4K20me1 and H4K20me2 are important in DNA replication and DNA repair, whereas H4K20me3 is a hallmark of silent heterochromatin. Studies have shown that H4K20me2 as the most abundant methylation state on histone H4, and H4K20me2 is widely distributed across the genome^10–12^, indicating its crucial roles in DNA-templated processes. Dysregulation of H4K20 methylation states leads to various diseases, including cancer^13^.

SUV420H1 is a member of the SUV4-20H family protein, responsible for catalyzing the majority of H4K20me2 and H4K20me3 modifications. This family consists of a unique N-terminal domain, a catalytic SET domain, and a Zn-binding post-SET domain. SUV420H1 mediated H4K20me2 has been linked to the recruitment of 53BP1 to DNA double-strand breaks (DSBs) regulating DNA damage repair^6, 8, 14^. H4K20me2 is also enriched at replication origins, and abrogating ORC1 recognition of H4K20me2 in cells impairs ORC1 occupancy at replication origins^7^. A recent study identified that histone variant H2A.Z promotes the catalytic activities of SUV420H1 on H4K20me2 mark deposition, which in turn facilitating ORC1 recruitment, regulating DNA replication^15^. Based on the catalogue of somatic mutations in cancer (COSMIC) database, somatic mutations of SUV420H1 are frequently occurred in different types of cancers^16^. In addition, it has shown that SUV420H1 dysregulation is associated with neurodevelopmental abnormalities^17–19^.

Several somatic cancer mutations in SUV420H1 exhibited lower enzymatic activity on nucleosomes^20^. While the structure of SUV420H1 is currently limited to the SET domain^21^. Recent years, significant advancements in the structures of major important histone methyltransferases in complex with nucleosomes providing vast insights into their enzymatic mechanisms and the relevance of diseases^22–25^. However, the specific mechanism by which SUV420H1 specifically recognizes and methylates histone H4 at lysine 20 (H4K20) at nucleosome level remains largely unclear. Additionally, the mechanism underlying the enhancement of SUV420H1 activities by H2A.Z-containing nucleosomes remains puzzling.

Here, we report the cryo-electron microscopy (cryo-EM) structures of SUV420H1 bound to H2A containing nucleosome (SUV420H1–NCP^H2A^) and H2A.Z containing nucleosome (SUV420H1–NCP^H2A.Z^) at an overall resolution of 3.58 Å and 3.68 Å, respectively. In combination with structure-guided functional experiments, our results provide an overview of how SUV420H1 interacts with the histones and DNA components of the nucleosome to specifically recognize and catalyze H4K20. Our results also elucidate how H2A.Z promoting SUV420H1 catalytic activity.

## Results

### Biochemical analysis of SUV420H1 and the overall structure of the SUV420H1– NCP^H2A.Z^ complex

The expression of full-length human SUV420H1 in *E. coli* cells turned out to be unstable for cryo-electron microscopy (cryo-EM) structural studies. Therefore, we screened SUV420H1 constructs with different truncations to improve SUV420H1 homogeneity. Finally, we found the construct that referred to as SUV420H1_1-390_ containing SUV420H1 residues 1-390 yielded high expression with ideal behavior in *E. coli* cells (hereafter SUV420H1 refers to SUV420H1_1-390_) (Fig. S1a). To stabilize the SUV420H1 and NCPs complex for cryo-EM studies., we have introduced H4K20M mutation and found that SUV420H1 bound to H4K20M NCPs (NCP^H4K20M^) more efficiency than canonical NCP^H4K20^ (Fig. S1b-c). Microscale thermophoresis (MST)-based binding analysis showed that SUV420H1 had around two-fold higher binding affinity for NCP^H4K20M^ than for canonical NCP^H4K20^ (Fig. S1c). Studies on H2A.Z facilitating the licensing and activation of replication origins revealed that H2A.Z-containing NCPs bind directly to the histone lysine methyltransferase SUV420H1, promoting H4K20me2 deposition^15^. As expected, MST-based binding analysis showed that SUV420H1 had around 3-fold higher binding affinity for NCP^H2A.Z^ than for NCP^H2A^ (Fig.1a). The end-point histone methyltransferase (HMT) assay verified that the enzymatic activity of SUV420H1 towards NCP^H2A.Z^ exhibited 1.6-fold as much as SUV420H1 towards NCP^H2A^ (Fig.1b). We then used Grafix^26^ method to further stabilize and purified the complex (Fig. S1d) and determined the cryo-EM structures of SUV420H1 in complex with H2A containing NCPs (SUV420H1–NCP^H2A^) and with H2A.Z containing NCPs (SUV420H1–NCP^H2A.Z^) at an overall resolution of 3.58 and 3.68Å, respectively (Fig.1c-d, Fig.S2, S3, Table S1). In both cryo-EM structures, SUV420H1 made extensive contacts with the DNA and histones of NCPs. More importantly, we observed additional density of SUV420H1 directly interacting with the H2A.Z-H2B acidic patch (Fig.1c-d).

### Nucleosome DNA and H4 tail recognition by SUV420H1

In both structures, Arg286 of SUV420H1 recognizes the phosphate backbone of one strand spanning the DNA minor groove at superhelix location 2 (SHL 2) (Fig. 2b). Arg286 is a human cancer associated mutational site^16^. To further consolidate its importance, we disrupted Arg286 with a mutation of R286A. It showed an 87% reduction in binding affinity with NCP compared to wild-type SUV420H1 and a loss of 46% catalytic activity towards nucleosome substrates (Fig. 2e-f), suggesting an important role of Arg286 in nucleosome recognition and H4K20 methylation.

**Figure 1.**
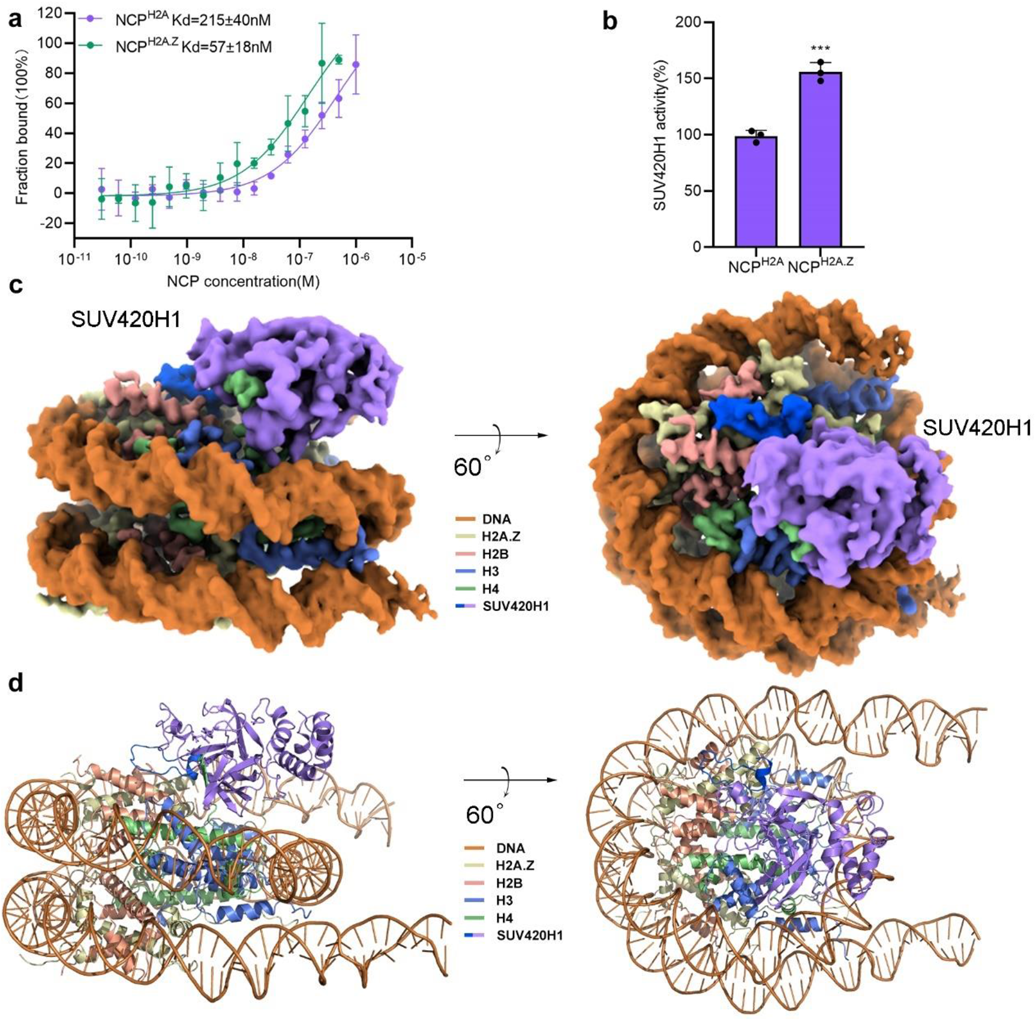
Biochemical analysis of SUV420H1 and the overall structure of the SUV420H1– NCP^H2A.Z^ complex. **a,** Microscale thermophoresis assays of the binding of human SUV420H1 to NCP^H2A^ and NCP^H2A.Z^. Dissociation constants (Kd) = 215 ± 40 nM (NCP^H2A^) and 57 ± 18 nM (NCP^H2A.Z^). Error bars represent mean ± SEM based on three independent measurements. **b,** Catalytic activity of the SUV420H1 on NCP^H2A^ and NCP^H2A.Z^ by end-point HMT assays *in vitro*. Each assay was repeated at least three times with similar results. n = 3 independent experiments, two-tailed, unpaired t-test. *** p=0.00086. **c, d,** Cryo-EM density map (**c**) and atomic model (**d**) of human SUV420H1–NCP^H2A.Z^ complex, shown in side (left) and top (right) view. The cryo-EM map is segmented and coloured according to the components of the SUV420H1–NCP^H2A.Z^ complex.

**Figure 2.**
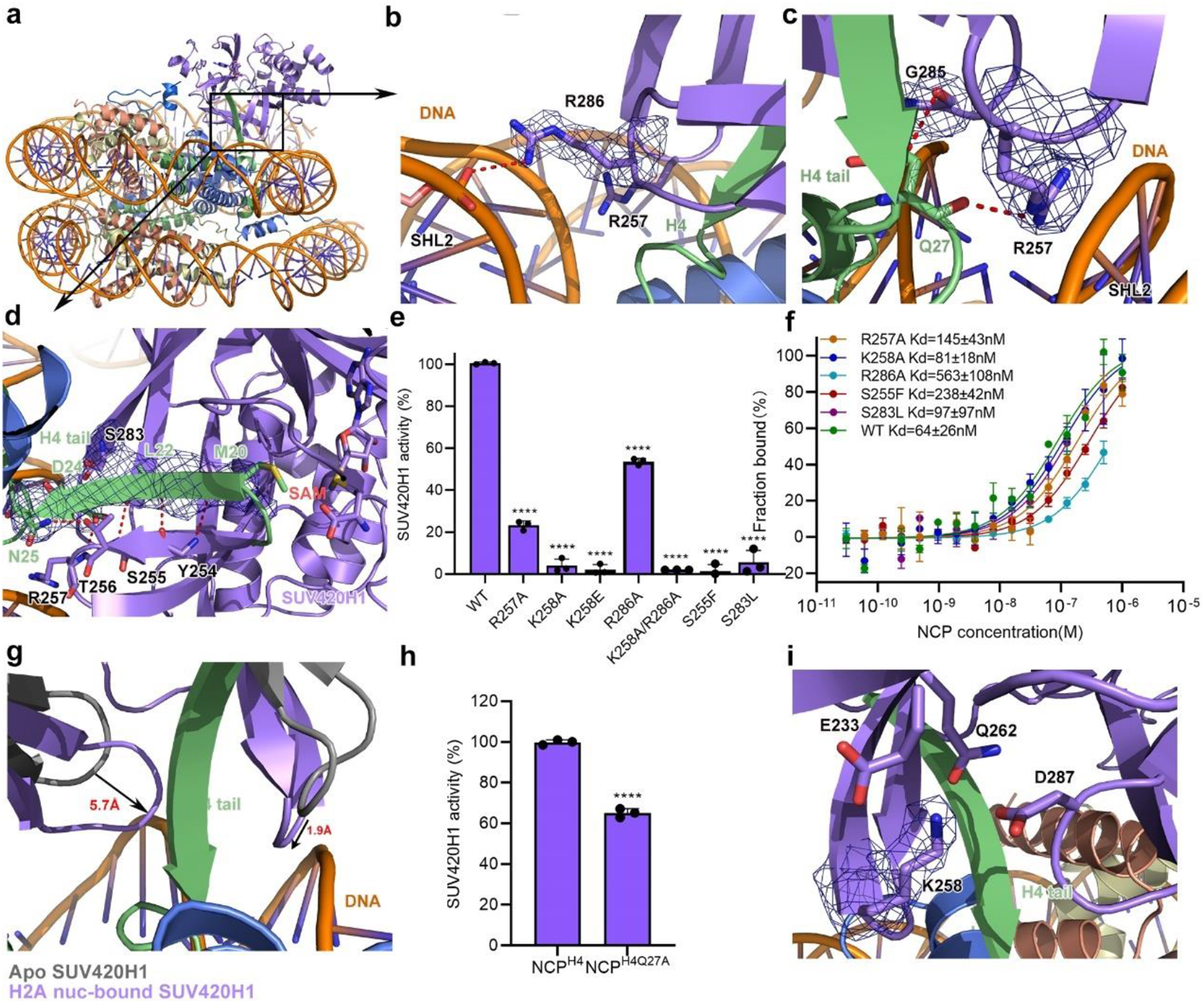
Interfaces between SUV420H1 and the H4 tail together with DNA. **a,** Overview of the recognition interfaces between SUV420H1 and nucleosomal H4 tail and DNA components. SUV420H1 domains are colored as in Fig. 1d. Histones H2A.Z and H2B are colored in yellow and pink, respectively. **b,** Detailed view of the interactions between SUV420H1 and the phosphate backbone of nucleosome SHL 2 DNA. Important residues at the interface are shown as sticks. **c,** Detailed view of the interactions between SUV420H1 and the H4 tail Q27. Important residues at the interface are shown as sticks. **d,** Detailed view of the recognition interfaces between SUV420H1 and nucleosomal H4 tail. Residues at the interface of H4 are shown as sticks. **e,** Catalytic activity of wild-type SUV420H1 and various mutants on NCP^H2A^ by end-point HMT assays in vitro. Adjusted p-values for pairwise ANCOVA comparison of wild type SUV420H1 and each mutant are reported: **** p<0.0001. **f,** MST binding assays of wild-type SUV420H1and SUV420H1 mutants on NCP^H2A^. Binding curves and Kd values are also shown. Error bars represent mean ± SEM based on three independent measurements. **g,** Alignment of the apo SUV420H1 (Protein Data Bank (PDB) code 3S8P, grey) with H2A NCP-bound SUV420H1 (colored as in Fig. 1d). Directions of shifted regions of H2A NCP-bound SUV420H1 are indicated with black arrows. **h,** Catalytic activity of SUV420H1 on NCP^H2A^ and NCP^H4Q27A^ by end-point HMT assays *in vitro*. Adjusted p-values for pairwise ANCOVA comparison of wild type SUV420H1 and each mutant are reported: **** p<0.0001. **i,** Detailed view of the interaction interface of SUV420H1 residue K258 in a negative charged pocket. The important residues are shown as sticks.

The positively charged side chain of Arg257 is pointed towards DNA minor groove through hydrogen bonds between Arg257 side-chain and main-chain of Gln27 of H4. The side-chain of H4 Gln27 also forms hydrogen bond with the main-chain of Gly285 of SUV420H1 (Fig. 2c). The side-chain of Lys258 inserts into a negative charged pocket including residues Glu233, Gln262 and Asp287 of SUV420H1 (Fig. 2i), further stabilizing the complex. Arg257, Lys258 and Asp287 are also mutational hotspots in human cancers^16^. Mutations of R257A and K258A showed 56% and 21% reduction in binding affinity compared to wild-type SUV420H1. R257A mutant showed a 77% catalytic activity loss towards nucleosomal substrates, whereas the K258A, K258E and K258A/R286A double mutant nearly lost the catalytic activity (Fig. 2e-f), suggesting that the catalytic activity of SUV420H1 is lost in the patients harboring these mutations. The mutation of Q27A in histone H4 leads to a 35% decrease in catalytic activity compared to wild-type SUV420H1–NCP^H2A^ complex (Fig. 2h).

The interactions between H4 tail and the *β* strand containing residues 252-256 of SUV420H1 are mainly mediated by hydrogen bonds between main-chain/side-chain atoms. Hydrogen bonds between Thr256 of SUV420H1 and Asn25, Asp24, Leu22 of H4, between Tyr254 of SUV420H1 and Leu22 of H4, between Tyr254 of SUV420H1 and Met20 of H4 tightly position the side-chain of Met20 of H4 facing towards the methyl donor of the ligand S-adenosylmethionine (SAM) through insertion into the hydrophobic catalytic pocket of SUV420H1 (Fig. 2d, S4a).

S255F and S283L are cancer associated mutations from the catalogue of somatic mutations in cancer (COSMIC) database^16^. In our structure, we found that S255 and S283 just locate at the entrance of that H4 tail insertion into the catalytic pocket (Fig. S4b). When the serine was mutated to phenylalanine or leucine, the bulky side chain will block the entry for H4 tail insertion. Thus, S255F and S283L mutations showed 73% and 34% loss in binding affinity, and a dramatically decreased in catalytic activity (Fig. 2e-f), which is consistent with a previous study ^20^.

Compared with the apo structure of SUV420H1 (PDB 3S8P)^21^, the NCP-bound activated SUV420H1 catalytic loops adopted a more compact conformation.

Specifically, the loop containing residue R286 undergoes a movement of approximately 5.7 Å towards the H4 tail. This conformational change brings R286 in closer proximity to the H4 tail, suggesting its involvement in the catalytic activity of SUV420H1. Similarly, the loop containing residue K258 moves around 1.9 Å towards the H4 tail. This movement indicates that K258 plays a role in the interaction with the H4 tail and may contribute to the catalytic activity of SUV420H1 when bound to the nucleosome core particles (Fig. 2g), further facilitating SUV420H1 recognition of nucleosome DNA and H4 tail. These observed conformational changes in the catalytic loops highlight their dynamic nature and their importance in mediating the recognition and binding of the H4 tail by SUV420H1 when it is in complex with nucleosomes. Interestingly, the structure of NCP-bound SUV420H1-H4 resembles the crystal structure of SUV420H2-H4 complex (PDB 4AU7), with the difference that the DNA interacting loops get closer to the nucleosomal DNA (Fig. S4c), illustrating the difference between the recognition of different substrates.

### C terminal domain of SUV420H1 recognizes H2A.Z-H2B acidic patch

Interestingly, in both of the cryo-EM density maps of SUV420H1–NCP^H2A^ and SUV420H1–NCP^H2A.Z^ complexes, we observed additional SUV420H1 density interacting with nucleosome H2A (H2A.Z)-H2B acidic patch that is good enough to be modeled. The residues 341-361 of SUV420H1 were built into this region by *de novo* modeling, assisted with the predicted models from AlphaFold Protein Structure Database^27^. Recognition of the acidic patch of the nucleosome by arginine anchors^28^ is widely found in histone lysine methyltransferases, including Dot1L, Set8 and COMPASS^24, 29, 30^. The structure reveals that two arginine anchors of the C terminal domain of SUV420H1 make extensive interactions with the acidic pocket of the nucleosome. The side chain of the positively charged residue Arg352 inserts into a negatively charged network that is composed of residues Glu64, Glu93 and Glu95 of H2A.Z, enabling SUV420H1 to engage with nucleosome by electrostatic interactions (Fig. 3b). Arg357 is positioned close to the negatively charged residues Glu64 and Glu67, forming electrostatic interactions (Fig. 3c). In addition to the arginine anchors, residue Tyr349 forms hydrogen bond with Glu59 in H2A.Z (Fig. 3f). To examine the functions of Tyr349, Arg352 and Arg357, we reconstituted Y349A, R352A, R357A mutations and measured the binding affinity and catalytic activity of the mutant proteins towards nucleosome *in vitro*. As expected, Y349A, R352A and R357A displayed 81%, 65% and 35% loss in binding dissociation constant (Kd) towards nucleosome substrates in comparison with wild-type SUV420H1 (Kd=64 nM) (Fig. 3d), respectively. Compared with the wild-type SUV420H1–NCP^H2A^ complex, the catalytic activity of the mutants dramatically decreased *in vitro* (Fig. 3e), suggesting that the interactions between residues Tyr349, Arg352 and Arg357 of SUV420H1 and the acidic patch of NCP are necessary for SUV420H1 to exhibit full methylation activity on NCP. Recurrent mutation of Arg357 have been identified in cancers^16^, we also reconstructed the cancer-associated mutation R357H and found the mutant displayed very low methylation activity, in support of the functional importance of the arginine anchor of SUV420H1 for maximal activity. The two arginine anchors and Tyr349 of SUV420H1 are highly conserved in SUV4-20H family proteins (Fig. S5), illustrating a common mechanism for nucleosome recognition by SUV4-20H family enzymes.

**Figure 3.**
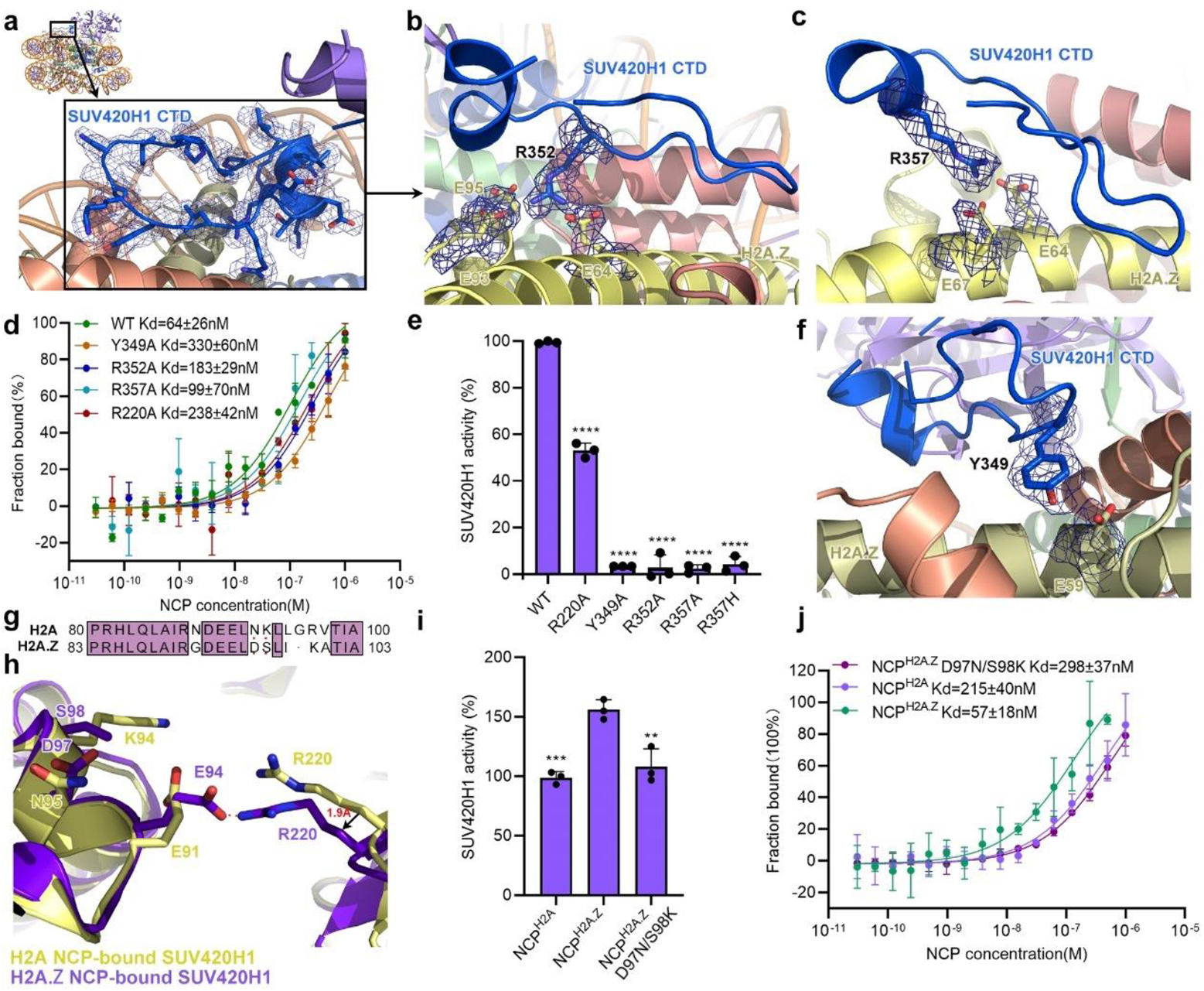
Contacts with H2A.Z-H2B acid path by CTD domain orient SUV420H1 onto nucleosome. **a,** Overview of the contacts between SUV420H1 CTD domain and nucleosomal H2A.Z-H2B acid path. SUV420H1 domains are colored as in Fig. 1d. Histones H2A.Z and H2B are colored in yellow and pink, respectively. **b, c, f,** Detailed view of the recognition of the acidic patch of the nucleosome by arginine anchors. Important residues at the interface are shown as sticks. **d,** MST binding assays of wild-type SUV420H1 and SUV420H1 mutants with NCP^H4M20^. Error bars represent mean ± SEM based on three independent measurements. Binding curves and Kd values are also shown. **e,** Catalytic activity of wild-type SUV420H1 and various mutants on NCP^H2A^ by end-point HMT assays in vitro. Each assay was repeated at least three times with similar results. N = 3 independent experiments, two-tailed, unpaired t-test. Adjusted p-values for pairwise ANCOVA comparison of wild type SUV420H1 and each mutant are reported: **** p<0.0001. **g,** Sequence alignment of the region containing D97 and S98 of H2A.Z and the corresponding region of H2A (in single-letter code). Same residues are boxed in purple background. **h,** Conformational changes and shifts of residue 220 in H2A NCP-bound SUV420H1 and H2A.Z NCP-bound SUV420H1. Directions of shifted regions are indicated with black arrows. **i,** Catalytic activity of SUV420H1 on NCP^H2A^, NCP^H2A.Z^ and NCP^H2A.Z^(D97N/S98K) by end-point HMT assays *in vitro*. n = 3 independent experiments, two-tailed, unpaired t-test. *** p=0.0006, ** p=0.0086. **j,** MST binding assays of wild-type SUV420H1 and SUV420H1 mutants on NCP^H2A^. Binding curves and Kd values are also shown.

The methylation activity of SUV420H1 towards H2A.Z mononucleosome substrates was greatly enhanced, comparing with H2A mononucleosome substrates as reported previously^15^, but this mechanism remains poorly understood. The different acidic patch residues D97/S98 in H2A.Z corresponding to N94/K95 in H2A are important in nucleosomal surface alteration further affecting chromatin factors binding^31^. We generated double mutant D97N/S98K of H2A.Z and measured the binding affinity and catalytic activity of SUV420H1 (Fig. 3g). SUV420H1 carried out comparable catalytic activity and binding affinity on H2A.Z D97N/S98K NCP to canonical NCP, which demonstrate that D97 and S98 play a dominant role in the enhanced activity of H2A.Z to SUV420H1 (Fig. 3i-j), in consistent with a previous study^15^. Next, we aligned SUV420H1–NCP^H2A^ complex with SUV420H1–NCP^H2A.Z^ complex, the two structures superimposed well with each other, except for some minor differences. Interestingly, we observed that residue R220 of SUV420H1 pointing to H2A/H2A.Z acidic patch undergoes slight conformational change, that the distance between R220 and E94 of H2A.Z is closer than that in SUV420H1–NCP^H2A^ complex (Fig. 3h, S6a-c). To further demonstrate the role of R220 in SUV420H1 methylation, we disrupted R220 with a mutation of R220A and measure the binding affinity and catalytic activity. Compared with the wild-type complex, the mutant displayed a 73% loss of binding affinity with NCP and a 47% decrease in catalytic activity, explaining the critical role of R220 in SUV420H1 methylation (Fig. 3d-e). Together, our research provides structural insights into the mechanism of H2A.Z enhancing SUV420H1 catalytic activity.

### Functional validation of SUV420H1–NCP complex

As we have observed multiple interactions between SUV420H1 and nucleosome that are critical for the binding and catalytic activity of SUV420H1 towards nucleosome substrate, we further tested the role of these interactions under physiological condition. We ectopically expressed SUV420H1_(Aa1-550)_ or SUV420H1_(Aa1-550)_ with a single K258E, S255F, S283L, Y349A, R352A, R357H or R220A mutation in a HeLa SUV420H1^−/−^ cell line. We found that compare to SUV420H1_(Aa1-550)_, mutants S255F and S283L show no rescue effect on both H4K20me2 and H4K20me3 (Fig. 4a), which is consistent with the critical role of Ser255 and Ser283 in the catalytic core of SUV420H1. We also found that mutants K258E, Y349A and R352A only show partial rescue effect on H4K20me2, and even less on H4K20me3 (Fig. 4a). Mutants R357H and R220A restored H4K20me2 almost as SUV420H1_(Aa1-550)_ did, but it cannot fully rescue H4K20me3 (Fig. 4a). These results supported the important role of Lys258, Tyr349, Arg352, Arg357 and Arg220 residues in facilitating the binding and catalytic activity on nucleosome substrate under physiological condition. We also noted that deletion of Aa1-62 of SUV420H1_(Aa1-550)_ did not impair its catalytic activity obviously (Fig. 4a), which is consistent with *in vitro* HMT assay (Fig. S8c). It has been reported that H4K20me2/3 is involved in the regulation of heterochromatin^6^. Consistently, through ATAC-seq we found that the number of open regions increased dramatically after SUV420H1 knock out (Fig. 4b). Then we quantify the dynamics of ATAC-seq signal in the peak regions. We found that 18222 regions are more opened in SUV420H1^−/−^ cells, while only 2846 regions are less open (Fig. 4c). Importantly, after ectopically express SUV420H1_(Aa1-550)_ in SUV420H1^−/−^ cells, 8857 (11.9%) of the peak regions significantly (p-value < 0.01) decreased ATAC-seq signal by two-fold (Fig. 4d), suggesting that these chromatin regions are restored to closed state. Through the same criteria, we analyzed the chromatin states by ATAC-seq after ectopically over-expressing SUV420H1_(Aa1-550)_ mutants in a HeLa SUV420H1^−/−^ cells. We found that, after over-expressing K258E, S255F, S283L, Y349A, R352A, R357H or R220A mutants, the number of peaks show decreased ATAC-seq signal is less than after over-expressing SUV420H1_(Aa1-550)_ (Fig. 4e), indicating that none of the mutants can restore the chromatin states as efficiently as SUV420H1_(Aa1-550)_ does. Through analyzing ATAC-seq signal in the 8857 “down” peaks as indicated in Fig. 4d, we found that the ATAC-seq signal decreased much less when over-expressing the mutants than over-expressing SUV420H1_(Aa1-550)_ (Fig. 4f). These results supported that K258E, S255F, S283L, Y349A, R352A, R357H and R220A mutation impaired the activity of SUV420H1 in restore the chromatin states, which is consistent with the defects of these mutants in rescuing H4K20me2/3.

**Figure 4.**
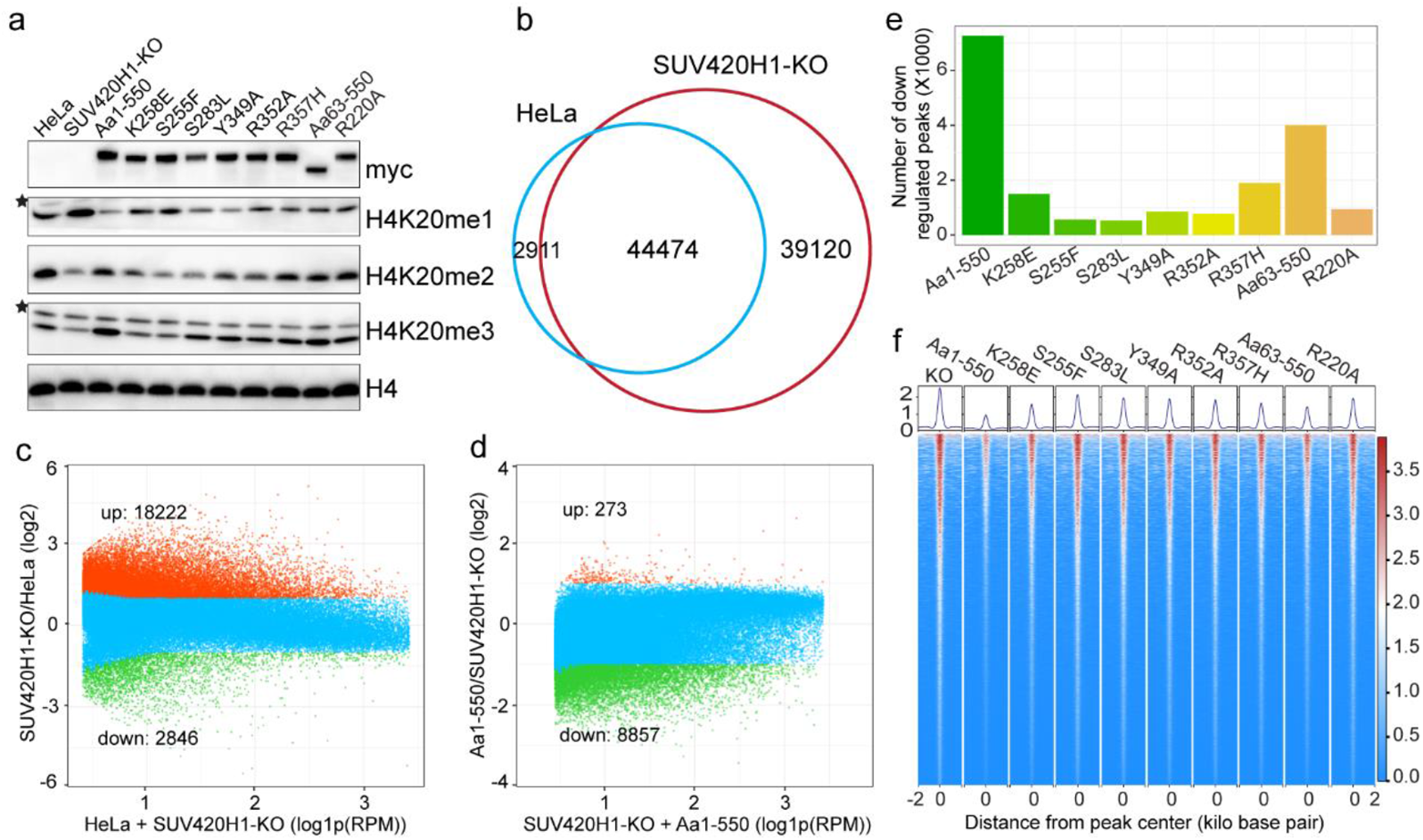
Functional validation of SUV420H1–NCP complex. **a,** Western blot shows the levels of H3K4me1/2/3 after over-expressing SUV420H1(Aa1-550) or SUV420H1(Aa1-550) with a single mutation of K258E, S255F, S283L, Y349A, R352A, R357H or R220A in SUV420H1^−/−^ HeLa cells, whose expression is indicated by myc. The stars indicate non-specific signal from H4K20me1 or H4K20me3 antibody. **b,** Venn plot shows the overlapping of ATAC-seq peaks from wild type and SUV420H1^−/−^ HeLa cells. **c,** Dot plot shows the dynamics of ATAC-seq signal in wild type and SUV420H1^−/−^ HeLa cells within the ATAC-seq peaks (n=75999) from SUV420H1^−/−^ HeLa cells. up (n=18222) and down (n=2846) indicate peaks with increased or decreased ATAC-seq signal filtered with 2-fold changes and p-value < 0.01. **d,** Dot plot shows the dynamic changes of ATAC-seq signal after over-expressing SUV420H1(Aa1-550) in SUV420H1^−/−^ HeLa cells, within the ATAC-seq peaks (n=75999) from SUV420H1^−/−^ HeLa cells. up (n=273), and down (n=8857) indicate peaks with increased or decreased ATAC-seq signal filtered with 2-fold changes and p-value < 0.01. **e,** Bar plot shows the number of peaks with decreased ATAC-seq signal after over-expressing SUV420H1(Aa1-550) or SUV420H1(Aa1-550) with a single mutation of K258E, S255F, S283L, Y349A, R352A, R357H or R220A, filtered with 2-fold changes and p-value < 0.01. **f,** Heatmap shows the ATAC-seq signal around the center of the 8857 “down” ATAC-seq peaks as indicated in Fig. 4d, from SUV420H1^−/−^ HeLa cells or SUV420H1^−/−^ HeLa cells over-expressing SUV420H1(Aa1-550) or SUV420H1(Aa1-550) with a single mutation as indicated in fig. 4e.

## Discussion

The direct interaction between the SUV420H1 and nucleosome demonstrates that SUV420H1 recognizes three elements, the H4 tail, the acidic patch and the DNA component to achieve a tight binding like other histone methyltransferases^24, 30, 32^. It is observed in our structure that somatic cancer-associated mutations S255F and S283L with larger sidechains block the insertion of the H4 tail into SUV420H1 SET domain at the entrance site (Fig. S4b), thus inhibiting the methylation activity of SUV420H1 confirmed in vitro and in vivo assay^20^. For cancer-associated mutation K258E, it disrupts the interaction of K258 with negative charged pocket (Fig. 2i), This disruption prevents the stabilization of the SUV420H1 complex bound to the nucleosome, leading to a loss of function. SUV420H1 anchors on nucleosome through R357 for its full activity. The cancer-related mutant R357H impairs the methylation activity of SUV420H1 *in vitro* and cannot fully rescue H4K20me3 in SUV420H1 in a HeLa SUV420H1^−/−^ cells. These findings provide structural evidence for how SUV420H1 recognizes and interacts with nucleosomes, and they elucidate the mechanisms through which disease-associated mutations in SUV420H1 inhibit its methylation activity in the context of nucleosomes.

A previous report has indicated that the DNA termini in H2A.Z nucleosomes are more flexible compared to H2A nucleosomes^33^ (Fig. S7c). From our research, there is a slight unwrapping of DNA in SUV420H1–NCP^H2A.Z^ complex structure compared to the H2A nucleosome (PDB 7KTQ) (Fig. S7b), while there isn’t DNA unwrapping of SUV420H1–NCP complex structure (Fig. S7a). These observations suggest that the flexibility of DNA termini in H2A.Z nucleosome leads to DNA unwrapping from the core histones in the SUV420H1–NCP^H2A.Z^ complex (Fig. S7d). It is important to note that these observations and speculations provide valuable insights into the structural dynamics and potential mechanisms underlying the interaction between SUV420H1 and H2A.Z nucleosomes. The increased flexibility of the DNA termini in H2A.Z nucleosomes could potentially facilitate the unwrapping of DNA upon SUV420H1 binding. This unwrapping event may have functional implications for the regulation of SUV420H1 activity and its specific targeting to H2A.Z-containing nucleosomes.

It has been reported that H4K20me2/3 are deposited at pericentric heterochromatin^10^ or telomeric heterochromatin^34^ by SUV420H1/2, which is recruited to these regions via Heterochromatin Protein 1 (HP1) in a H3K9me3 dependent manner^10^. In skeletal muscle stem cells (MuSCs), SUV420H1 promotes the formation of facultative heterochromatin to regulate MuSC quiescence^35^. However, whether and how H4K20me2/3 is directly involved in the regulation of heterochromatin at genome-wide is not clear. In our study, we found by ATAC-seq, which is widely used to map open chromatin regions, that after knock out of SUV420H1 in HeLa cells, the chromatin transit into a hyper-open state, suggesting that SUV420H1 mediated H4K20me2/3 may also play a role in coordinating the activity of *cis*-regulatory elements. In concordance with this speculation, SUV420H1 catalyzed H4K20me2 can recruit ORC1 to facilitate the activation of replication origins^15, 36^. Importantly, we identified critical interactions within the SUV420H1-NCP complex that are required for the full activity for SUV420H1, which may be useful to dissect the role of H4K20me2/3 in the regulation of heterochromatin without disrupt SUV420H1 entirely.

We observed an additional density in our cryo-EM density map that could not be attributed to a specific atom model, possibly corresponding to the N-terminal region of SUV420H1 (Fig. S8a). This observation led us further investigate the binding affinity and enzymatic activity of different SUV420H1 truncations. Interestingly, we found that SUV420H1_(1-335)_ (Kd =68 nM), exhibited comparable binding affinity to nucleosomes as SUV420H1_(1-390)_ (Kd =68 nM). However, the truncation SUV420H1_(63-390)_, which lacks residues 1-62 of the N-terminal region, displayed a five-fold decrease in binding affinity (Kd = 338 nM) compared to SUV420H1_(1-390)_ (Fig. S8b). Despite this decrease in binding affinity, SUV420H1_(63-390)_ demonstrated comparable enzymatic activity to SUV420H1_(1-390)_ (Fig. S8c), indicating that the deletion of residues 1-62 primarily affects the binding affinity of SUV420H1 to nucleosomes without impairing its methylation activity. Additionally, your results suggest that the C-terminal domain (CTD) of SUV420H1 is essential for its methylation activity towards nucleosome substrates.

Overall, our findings provide valuable structural insights into the mechanisms of nucleosome recognition and methylation by SUV420H1. Furthermore, we identified key residues and interaction interfaces that contribute to the enhanced activity of SUV420H1 towards nucleosomes containing the variant histone H2A.Z compared to canonical nucleosomes. These structural elements and interactions uncovered in our study serve as important targets for further functional investigations of SUV420H1 and its role in chromatin regulation.

## Material and Methods

### Protein expression and purification

Human SUV420H1 protein, its truncations or mutants were cloned into a modified pET28a vector with an N-terminal 6×His-TEV tag. The expression and purification of SUV420H1 and mutants were performed using the same protocol. In general, SUV420H1 was transformed into *Escherichia coli* BL21(DE3)-RIL competent cells and were induced with 0.4 mM IPTG at 18 °C for an overnight expression. The cells were centrifuged at 5,000 rpm for 15 min at 4 °C, resuspended in lysis buffer B (25mM HEPES pH7.5, 500mM NaCl, 5% glycerol, 1 mM TCEP) and then sonicated for around 30 min. The supernatant was collected by centrifugation of the cell lysate at 20,000 rpm for 1 h and was added to Ni-NTA resin (BOGELONG) that had been equilibrated in buffer A (25mM HEPES pH 7.5, 500 mM NaCl, 5% glycerol, 1 mM TCEP). The beads were incubated with the lysate at 4°C for 1 h, poured into a gravity flow column and washed with buffer A. Protein was eluted using buffer B (25mM HEPES pH7.5, 500mM NaCl, 5% glycerol, 1 mM TCEP, 300 mM imidazole). After elution 1 mg of TEV protease was added directly to the eluate and dialyzed overnight at 4°C against SUV420H1 dialysis buffer (25 mM HEPES pH 7.5, 500 mM NaCl, 5% glycerol, 1 mM DTT). Precipitate was removed by centrifugation and filtration through a Amicon Ultra spin concentrator (Millipore). To separate SUV420H1 from TEV and the cleaved 6×His tag, the sample was again loaded onto the Ni-NTA resin equilibrated in buffer A, collecting the flow through. The protein was concentrated and was further purified by a Superdex increase 200 16/600 size exclusion chromatography column (GE Healthcare) that was pre-equilibrated with SEC buffer (25 mM HEPES pH 7.5, 300 mM NaCl, 5% glycerol, 1 mM DTT). Peak fractions corresponding to SUV420H1 were pooled and the pure concentrated protein (15 mg/ml) was stored at −80°C freezer until use.

### Nucleosome reconstitution

Histone H2A, H2A.Z, H2B, H3.1, H4 were cloned into pET22b vector. H4K20M was generated using site-mutagenesis introduced by primers.

The expression and purification of histones were performed using the same protocol. Briefly, the histone plasmid was transformed into *E. coli* BL21(DE3) and expressed in an insoluble form at 37℃. The cells were lysed and the insoluble bodies were collected, dialyzed in the denaturing buffer (20 mM sodium acetate pH 5.2, 7 M urea, 0.2–1 M NaCl, 1 mM EDTA and 5 mM 2-mercaptoethanol). After purified by a SP sepharose HP (GE Healthcare) column, the histone proteins were dialyzed completely against distilled water containing 2 mM β-mercaptoethanol, and were concentrated to 2mg/ml and then lyophilized to store at −80 °C.

The plasmid containing 12×167-bp 601 DNA fragment^37^ was constructed using *E. coli* DH5α strain. The pUC19-12×167 plasmid was large-scale purified and the 167-bp fragment was excised from plasmid by EcoRV digestion and isolated from the digestive product as described^38^.

Nucleosome core particles were reconstituted as described previously^39, 40^. In brief, histone octamers were prepared by mixing H2A, H2B, H3.1 and H4 in refolding buffer containing 2 M NaCl. Histone octamers were isolated by size-exclusion chromatography, concentrated and stored at −80 °C. Then, the purified histone octamer was mixed with the 167-bp 601 DNA at a molar ratio of 1:1.1 and was dialysed against reconstitution buffer (10 mM Tris-HCl pH 7.5, 1 mM EDTA, 1 mM DTT and 0.25–2 M KCl) by salt gradient dilution for 36 h. The reconstituted NCP was further purified through superdex increase 200 16/600 gel filtration, and was then stored at 4 °C.

### Electrophoretic mobility shift assay

SUV420H1 was mixed with 1 pM NCP^H2A^ or NCP^H2A.Z^ at a molar ratio of 0:1, 5:1, 9:1 and 13:1 to a total volume of 10 μl. After incubation on ice for 30 min in electrophoretic mobility shift assay (EMSA) buffer (25 mM HEPES pH 7.5, 50 mM NaCl, 50 mM KCl, 2 mM DTT, 40 μM SAM), the samples were loaded onto a 6% native TBE gel at 4 °C. Electrophoresis was performed at 4 °C for 90 min at a constant voltage of 120 V. The resulting gels were visualized by GelRed staining and visualized.

### Assembly of the SUV420H1–NCP complexes

Wild-type SUV420H1 was mixed with nucleosome containing H2A or H2A.Z at a molar ratio of 10:1 in binding buffer containing 50 mM KCl, 50 mM NaCl, 25 mM HEPES pH 7.5, 40 μM SAM and 2 mM DTT at 4 °C for 15 min.

The above samples were then subjected to GraFix^26^. Five hundred microliters of sample were put into the 12.5-ml tubes with a 10–30% linear glycerol gradient and a 0–0.025% linear glutaraldehyde gradient in 25 mM HEPES pH 7.5, 100 mM NaCl and 1mM DTT, then centrifugated at 35,000 rpm for 16 h using a Beckman SW41Ti rotor at 4 °C. Following centrifugation, the sample was divided into 200 μL each fraction from top to bottom, and the fractions were examined by 6% native TBE gel and negative-staining EM. Fractions containing uniform, properly sized particles were pooled and centrifugated to remove glycerol and concentrate.

### Cryo-EM sample preparation and data collection

For the cryo-EM specimen preparation of both the SUV420H1–NCP^H2A^ and the SUV420H1–NCP^H2A.Z^ complexes, Quantifoil R1.2/1.3 200 mesh Au grids were glow-discharged for 40 s using a Gatan Plasma System. With the Vitrobot (Thermo Fisher Scientific) set to 8 °C, 100% humidity, 3 μl of the samples were loaded onto the grid waiting for 10 s and then blotted for 3.0 s, directly plunged into liquid ethane.

Micrographs were collected on a Titan Krios microscope, operated at 300 kV, equipped with a K3 Summit direct electron detector (Gatan). SerialEM software (Thermo Fisher Scientific) was used for automated data collection^41^ in super-resolution mode at a nominal magnification of 130,000×, corresponding to a calibrated pixel size of 0.92 Å at the object scale. The defocus range was set from −1 μm to −2 μm. Each micrograph was dose-fractioned to 32 frames at a dose rate of 1.5652 e^−^/pixel/s, with a total exposure time of 2.1 s, resulting in a total dose of about 50 e^−^/Å^2^.

### Image processing

Both the SUV420H1–NCP^H2A^ and SUV420H1–NCP^H2A.Z^ data sets were processed in RELION 3.0^42^ and CryoSPARC ^43^. Motion correction and dose-weighted motion correction were performed using MotionCor2^44^. Gctf was used for CTF parameter estimation^45^. For the SUV420H1–NCP^H2A^ data set (4,106 micrographs), 1,201,595 particles were automatically picked and extracted in RELION. The autopicked particles were subjected to 4 rounds of 2D classifications. Then, 873,129 selected particles from the last 2D classification were subjected to 3D classification. The class of 158,987 particles with good SUV420H1 density were selected and re-extracted for a round of refinement. After that, to obtain better SUV420H1 density, local classification procedures were performed with the “--skip_alignment” option in RELION, and a few critical parameters, such as the mask size and regularization parameter T, were extensively tested. Finally, a total of 35,448 particles were selected for further refinement and postprocessing, yielding final reconstructions at overall resolution of 3.68 Å. The final selected good particles were transferred to CryoSPARC for a second refinement with some regions with better density.

For the SUV420H1–NCP^H2A.Z^ data set (6,794 micrographs), the processing procedures were the same. 1,782,793 particles were automatically picked and subjected to 3 rounds of 2D classifications in RELION. The selected 984,971 particles were subjected to two rounds of 3D classifications. Then 308,576 particles were selected and re-extracted for a round of refinement. Local classification procedures were performed with the “--skip_alignment” option in RELION, with the mask size and regularization parameter T tested. Then 45,377 particles were selected for final refinement and postprocessing, yielding an overall resolution of 3.68 Å. The final selected good particles were transferred to CryoSPARC for refinement and get a map with better SUV420H1 density.

### Model building, refinement and validation

Cryo-EM structure of NCP complex (PDB: 5WCU)^46^and crystal structures of the SET domain of human SUV420H1 (PDB: 3S8P)^21^ were used as initial models. These initial models were docked into the cryo-EM density maps in Chimera^47^ and manually adjusted in Coot^48^. The residues 341-361 of SUV420H1 were manually built *de novo* in Coot with the help of the predicted models from AlphaFold Protein Structure Database^27^. The atomic models were further refined against the density maps using Phenix.real_space_refine^49^ with the application of secondary structure restraints, geometry restraints and DNA-specific restraints. The quality of the final atomic models was evaluated by MolProbity^50^. Chimera, ChimeraX^51^ and PyMOL (http://pymol.org) were used for figure preparation.

### HMT assays

For a 20-μl end-point HMT reaction, 500 nM wild-type or mutants, 40 μM SAM, and 2 μM NCP^H2A^ or NCP^H2A.Z^ were mixed in the buffer of 25 mM HEPES-NaOH pH 7.5, 50 mM NaCl, 50 mM KCl, 5mM MgCl_2_, 0.1 mg/ml BSA and 1 mM DTT, and were incubated at 32°C for 15 min or 3 min. The reaction was stopped by adding 4 μl of 0.5% trifluoroacetic acid (TFA), and the HMT activity was evaluated using an MTase-Glo Methyltransferases Assay Kit (Promega). The luminescent signal that corresponds to the production of SAH was measured using synergy H1 microplate reader in a white 96-well plate. Each reaction was run in triplicate, and is reported as the mean ± s.d.

### MST assays

All MST-based experiments were performed on a Monolith NT115 Pico RED machine with 20% MST power and medium LED power. All of the proteins used in this assay have a 6×His-TEV tag at the N terminus. A Monolith His-tag Labeling Kit RED-tris-NTA (Nano Temper) was used to label the His-tagged proteins. For each reaction, 50 nM protein and 10 nM fluorescent dye were dissolved in 120 μL buffer containing 50 mM NaCl, 20 mM HEPES pH 7.5, 50 mM NaCl, 50 mM KCl, 1 mg/ml BSA, 0.05% Tween-20, 40 μM SAM and 2 mM DTT, and incubated at room temperature for 30 min. The nucleosomes were serially diluted in PCR tubes from 2 μM to 15.25 pM, then equal volumes of labelled proteins were added. After incubating for another 15 min, the samples were transferred into capillaries (Monolith NT115 Standard Treated Capillaries, MO-K002) in sequence. Dissociation constants (Kd) were fitted with the MO Affinity Analysis software. Each reaction was run in triplicate.

### Western Blot

Whole cell extract samples were prepared by extracting cell pellet with Nuclei Lysis Buffer (10 mM Tris-HCl, pH 7.5, 10 mM NaCl, 0.1% SDS) with PMSF. The antibody used were anti-H4K20me1 (Abcam, ab9051), anti-H4K20me2 (Abcam, ab9052), anti-H4K20me3 (Abcam, ab9053), anti-myc (Cell Signaling Technology, 2276S), anti-H4 (Cell Signaling Technology, 14149S).

### Assay for transposase accessible chromatin using sequencing (ATAC-seq)

ATAC-seq was performed according to Buenrostro et al.^52^ with minor modifications. Briefly, nuclei were extracted in Nuclei Buffer (10 mM Tris-HCl, pH 7.5, 10 mM NaCl, 3 mM MgCl_2,_ 0.1% Triton X-100) with Roche Complete Protease Inhibitor EDTA-Free. After counting nuclei, 1×10^5^ nuclei were incubated with 50 µL transposition mix containing 2 µL Tn5 transposase (Vazyme, TD711) at 37 °C for 30 min. Afterwards, 50 µL 2xStop buffer (20 mM EDTA, 0.2% SDS, 200 ug/ml Proteinase-K) were added to stop the reaction. The mixture were further incubated at 55 °C for 30 min, the DNA were extracted with Phenol:Chloroform:Isoamyl Alcohol (25:24:1), and precipitated by cold ethanol. 25 µL Tris buffer (10 mM Tris-HCl, pH 7.5) were used to resolve the DNA pellet, and 11.5 µL were used for Nextera library amplification. The library was purified with 1X AMPure XP beads (Beckman Coulter, A63881), and sequenced on DNBSEQ-T7 platform with PE150 mode.

### Sequencing data analysis

Paired-end reads were trimmed for adaptor sequence using cutadapt v4.3^53^ with parameters: -a CTGTCTCTTATACACATCT -A CTGTCTCTTATACACATCT -e 0.1 - n 2 -m 35 -q 30 –pairfilter = any, and then mapped to hg38 using Bowtie2 v2.5.1^54^ with parameters: −I 10 -X 1000 −3 5 --local --no-mixed --no-discordant --no-unal. Duplicates were marked using picard MarkDuplicates v2.27.5 (https://broadinstitute.github.io/picard/) with default parameters and removed using samtools view v1.17^55^ with parameters: -f 2 -F 1024 -q 10. ATAC peaks were detected with macs2 callpeak v2.2.7.1^56^ with parameter -f BAMPE -m 10 50. Signals within peaks were annotated using multiBigwigSummary, and heatmap was generated using plotHeatmap from deeptools v3.5.2^57^.

## Acknowledgments

We thank the cryo-EM facilities of Southern University of Science and Technology for technique assistance. We also thank Ning Gao and Ningning Li at Peking University for helpful discussion. This work was supported by the National Natural Science Foundation of China (32200473) and Shenzhen Science and Technology Program grant (RCYX20210706092045078 and 20220815154711001) to WL. This work was also supported by the National Natural Science Foundation of China (32000423) and Shenzhen Science and Technology Program grant (20220817134430001) to ZW.

## Author contributions

W.L. designed and supervised the research; F.L. reconstituted the complexes, performed cryo-EM sample preparation, data collection, processing and model building under the supervision of W.L.; F.L. performed biochemical analysis with the help of W.S., C.L., Y.Z., and M.M.; Z.W. and R.Z. performed rescue experiments in SUV420H1^−/−^ cells, Z.W. performed bioinformatic analysis; W.L. and Z.W. prepared the manuscript with contributions from all authors.

## Data and materials availability

Cryo-EM maps of SUV420H1-NCP^H2A^ and SUV420H1-NCP^H2AZ^ have been deposited in the Electron Microscopy Data Bank (EMDB) with accession codes EMD-36265 and EMD-36264, respectively. Atomic coordinates have been deposited in the Protein Data Bank (PDB) with accession codes 8JHG and 8JHF. High throughput sequencing data generated in this study were deposited to Gene Expression Omnibus (GEO) under accession XXXX. All other data are available from the corresponding author upon reasonable request.

