## Supplemental information for "Structural basis of nucleosomal H4K20 recognition and methylation by SUV420H1 methyltransferase"

### Supplementary Information

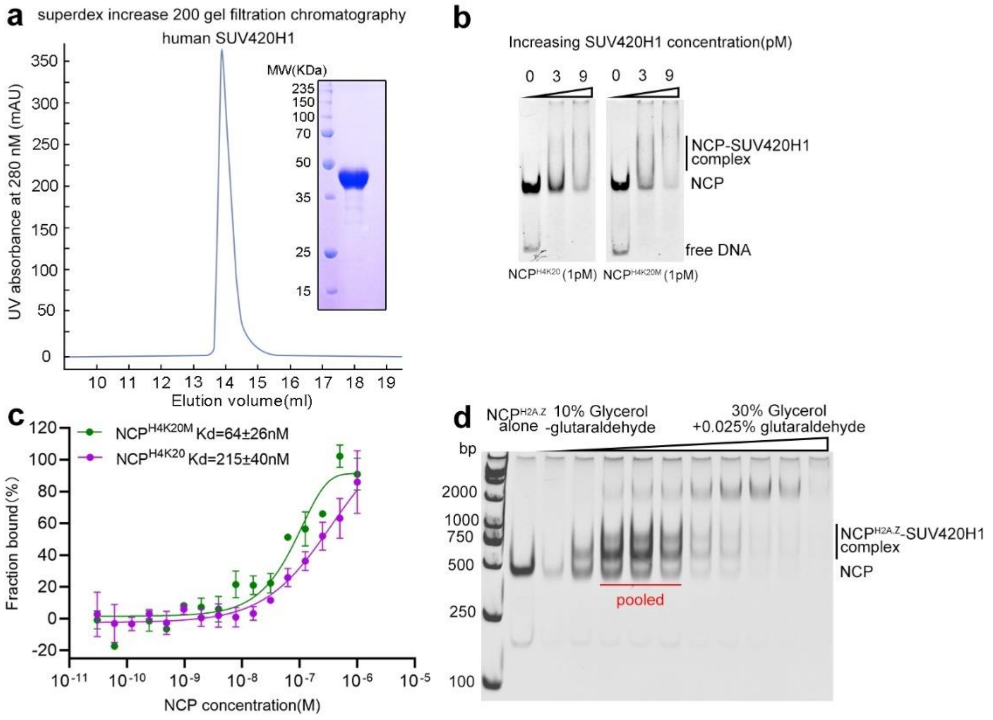

**Figure S1. Purification of SUV420H1, EMSA analysis and GraFix fractionation of SUV420H1 in complex with NCP. a,** Gel filtration and SDS–PAGE analysis of the human SUV420H1 (residues 1–390). Experiments were repeated at least three times with similar results. **b,** EMSA analysis of SUV420H1 binding to $NCP^{H4K20}$ and $NCP^{H4K20M}$. The concentrations of added SUV420H1 are indicated above the lanes. **c,** MST binding assays of wild-type SUV420H1 on $NCP^{H4K20}$ ( Kd=215±40nM ) and $NCP^{H4K20M}$ ( Kd=64±26nM ). Binding curves and Kd values are also shown. Error bars represent mean ± SEM based on three independent measurements. **d,** Representative images of 6% Native PAGE, stained with ethidium bromide after GraFix fractionation. Fractions containing cross-linked species indicative of a protein complex were used in structural studies.

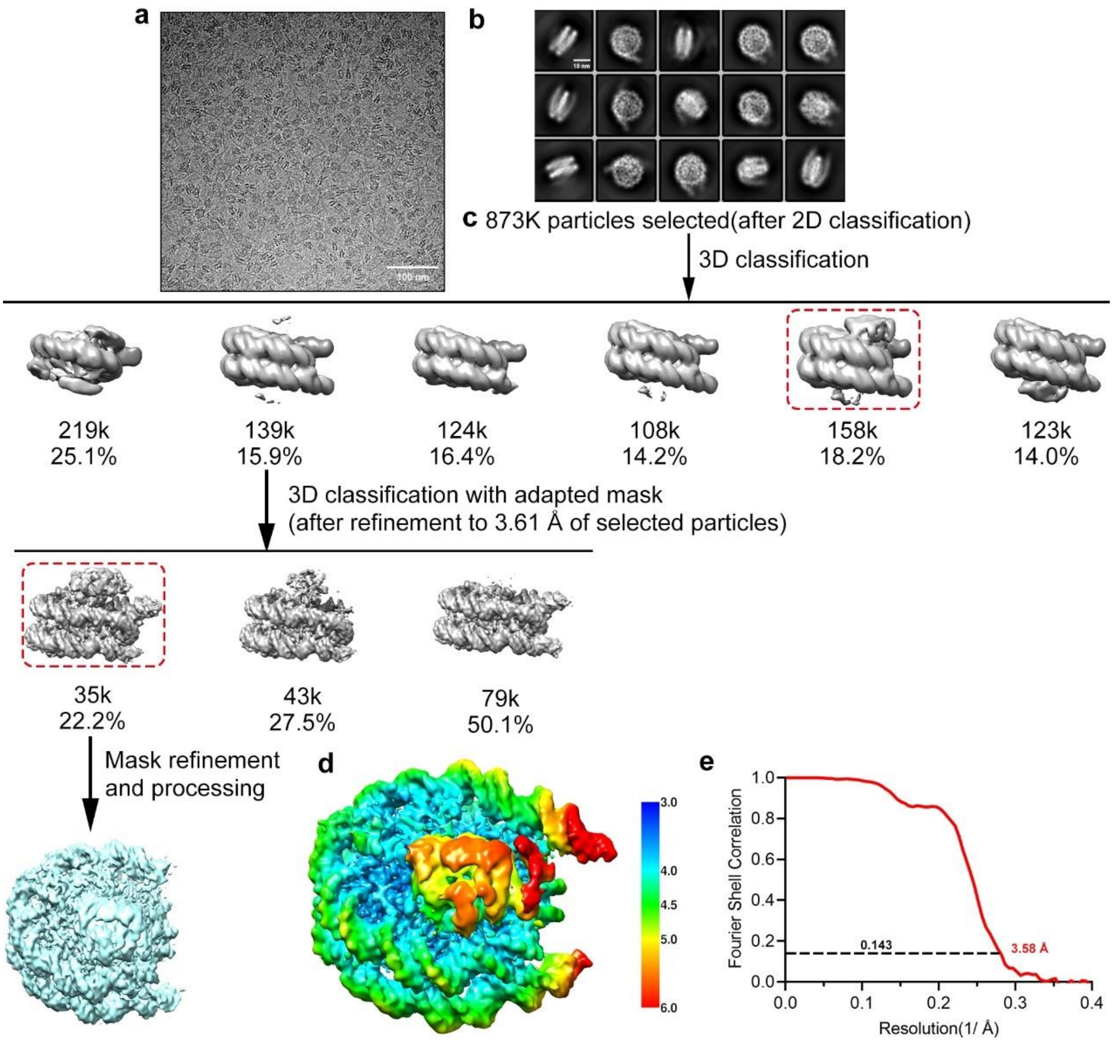

**Figure S2. Cryo-EM structural analysis of SUV420H1 in complex with NCP$^{H2A}$. a,** Representative cryo-EM micrograph from a total of 4,106 micrographs of the SUV420H1–NCP$^{H2A}$ complex (low-pass-filtered to 20 Å). Scale bar, 100 nm. **b,** Selected 2D class averages of the SUV420H1–NCP$^{H2A}$ complex. Scale bar, 10 nm. Box size 216, pixel size 0.92 Å. **c,** Workflow of the cryo-EM data processing procedures for the SUV420H1–NCP$^{H2A}$ complex. It includes several rounds of 2D and 3D classification, refinement and masked refinement. **d,** Local-resolution map of the SUV420H1–NCP$^{H2A}$ complex final density map. **e,** FSC curve of the SUV420H1–NCP$^{H2A}$ complex final density map.

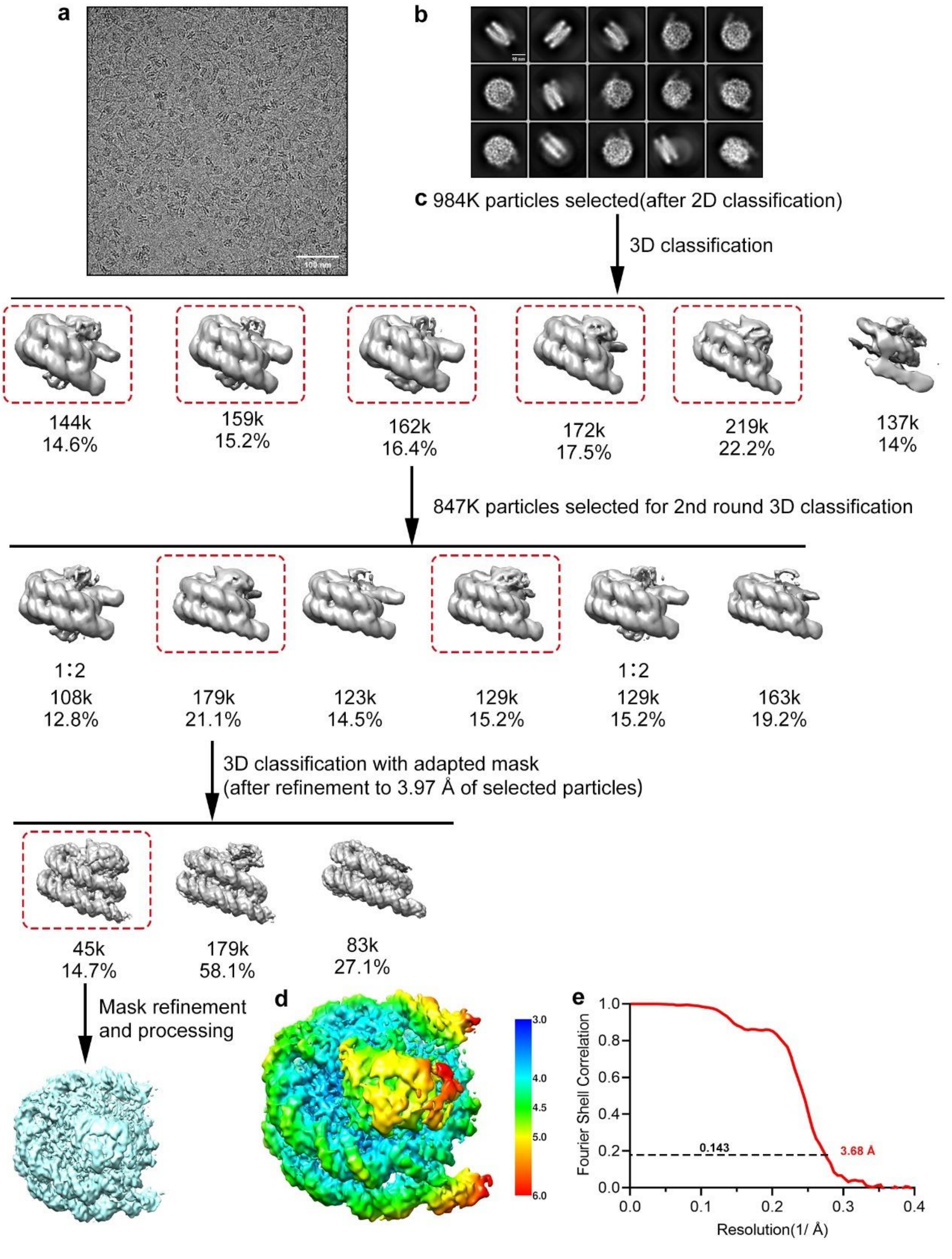

**Figure S3. Cryo-EM structural analysis of SUV420H1 in complex with $NCP^{H2A.Z}$. a,** Representative cryo-EM micrograph from a total of 6,915 micrographs of the SUV420H1–$NCP^{H2A.Z}$ complex (low-pass-filtered to 20 Å). Scale bar, 100 nm. **b,** Selected 2D class averages of the SUV420H1–$NCP^{H2A.Z}$ complex. Scale bar, 10 nm. Box size 216, pixel size 0.92 Å. **c,** Workflow of the cryo-EM data processing procedures for the SUV420H1–$NCP^{H2A.Z}$ complex. It includes several rounds of 2D and 3D classification, refinement and masked refinement. **d,** Local-resolution map of the SUV420H1–NCP$^{H2A.Z}$ complex final density map. **e,** FSC curve of the SUV420H1–NCP$^{H2A.Z}$ complex final density map.

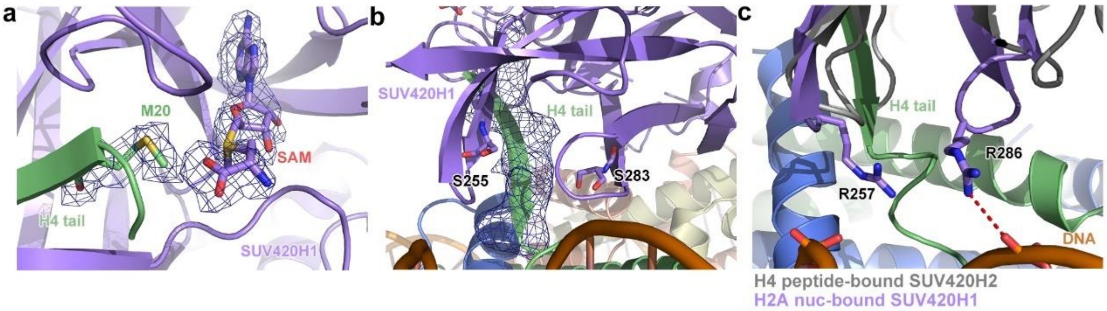

**Figure S4. Cryo-EM density maps. a,** The active site of SUV420H1$^{SET}$ within the SUV420H1–NCP complex, shown with the density maps of the cofactor-product SAM and the substrate residue H4K20 in stereo mode. **b,** Detailed view of the location of SUV420H1 S255 residue and S283 residue. The residues are shown as sticks and the cryo-EM density of the H4 tail residues 20-25 are shown in mesh. **c,** Alignment of the SUV420H2-H4peptide structure (PDB:4au7, shown in grey) with SUV420H1–NCP$^{H2A}$ complex structure.

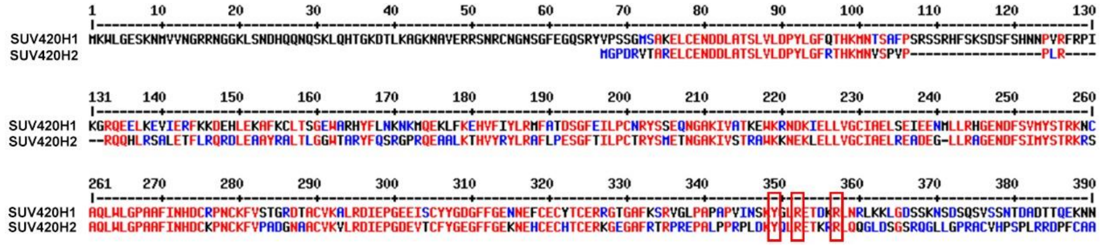

**Figure S5. Sequence alignment of SUV420H1 and SUV420H2. The two arginine anchors and Tyr349 residues are framed in red.**

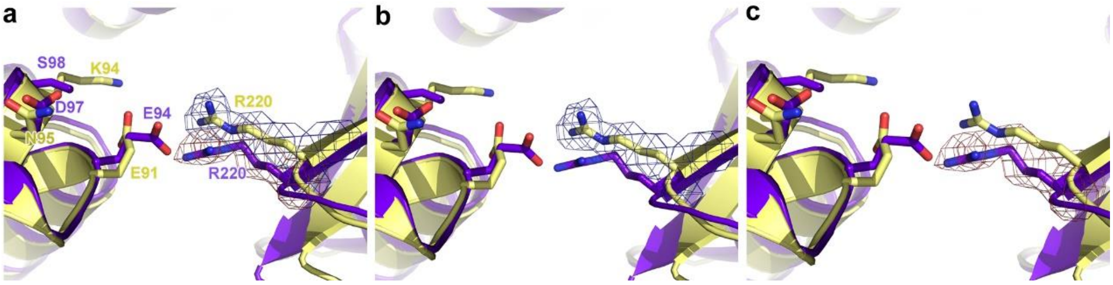

**Figure S6. Detailed view of the electron microscopy density maps of R220 in SUV420H1-NCP$^{H2A}$ and SUV420H1-NCP$^{H2A.Z}$ complexes. a,** Alignment of SUV420H1-NCP$^{H2A}$ and SUV420H1-NCP$^{H2A.Z}$ complexes showing the area comprised of R220 in SUV420H1 and acidic patch in NCP. SUV420H1-NCP$^{H2A}$ is colored in yellow, SUV420H1-NCP$^{H2A.Z}$ is colored in purple. **b,** The cryo-EM density of R220 in SUV420H1-NCP$^{H2A}$ complex is shown in mesh and contoured at 2σ level. **c,** The density map of R220 in SUV420H1-NCP$^{H2A.Z}$ complex is shown in mesh and contoured at 2.5σ level.

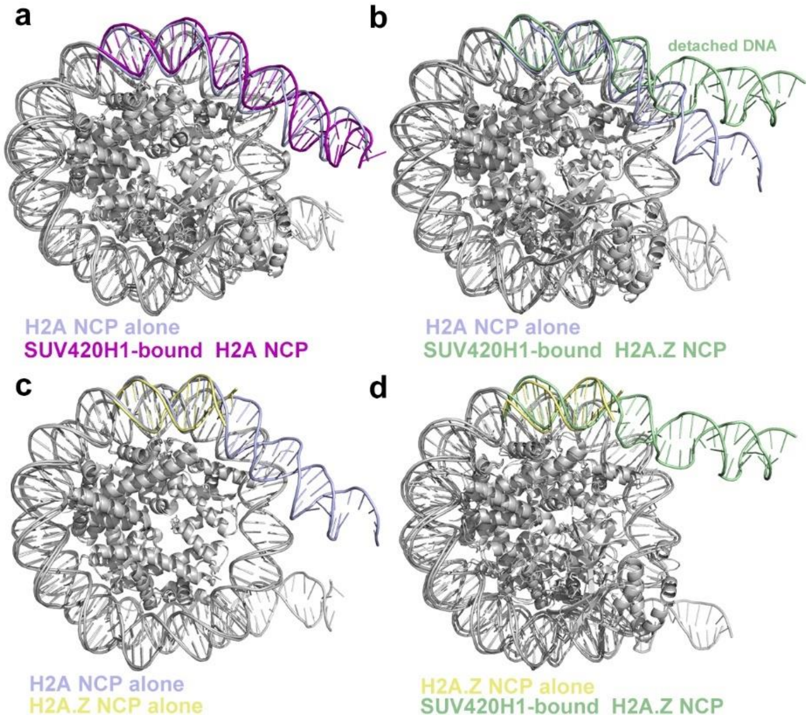

**Figure S7. Overlapped nucleosome structures of H2A/H2A.Z alone nucleosomes and SUV420H1-bound H2A/H2A.Z nucleosomes. a,** Overlapped nucleosome structures of H2A alone nucleosome (PDB 7KTQ) and SUV420H1-bound H2A nucleosome. The DNA at SHL 6 and SHL 7 are colored in blue (H2A alone NCP) and purple (SUV420H1-bound H2A NCP), respectively. **b,** Overlapped nucleosome structures of H2A alone nucleosome and SUV420H1-bound H2A.Z nucleosome. The DNA at SHL 6 and SHL 7 are colored in blue (H2A alone NCP) and green (SUV420H1-bound H2A.Z NCP), respectively. **c,** Overlapped nucleosome structures of H2A alone nucleosome and H2A.Z alone NCP nucleosome (PDB 7MIX). The DNA at SHL 6 and SHL 7 are colored in blue (H2A alone NCP) and yellow (H2A.Z alone NCP), respectively. **d,** Overlapped nucleosome structures of H2A.Z alone nucleosome and SUV420H1-bound H2A nucleosome. The DNA at SHL 6 and SHL 7 are colored in yellow (H2A.Z alone NCP) and green (SUV420H1-bound H2A.Z NCP), respectively.

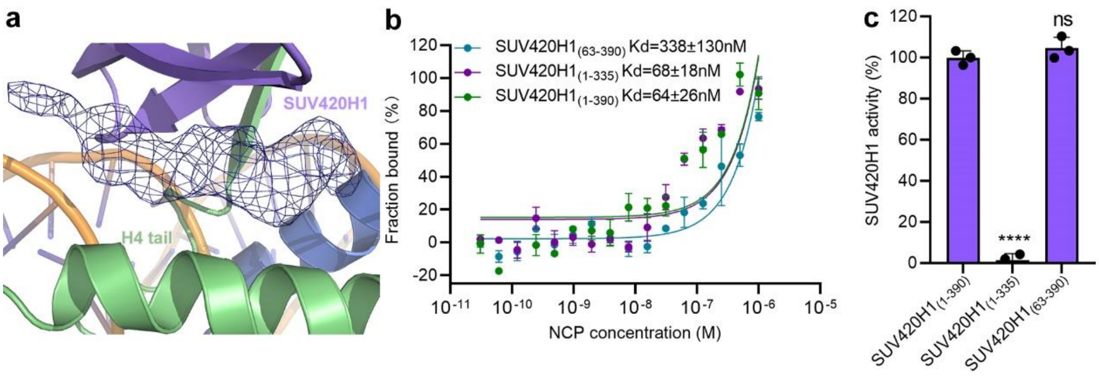

**Figure S8. Cryo-EM density map and biochemical assays of SUV420H1 truncations on nucleosomes *in vitro*. a,** The unmodeled additional density in SUV420H1-bound H2A nucleosome as shown in mesh. **b,** MST assays of the binding to $NCP^{H4K20M}$ of wild-type $SUV420H1_{(1\text{-}390)}$ (Kd=64±26nM), $SUV420H1_{(1\text{-}335)}$ (Kd=68±18nM) and $SUV420H1_{(63\text{-}390)}$ (Kd=338±130nM). Binding curves and Kd values are also shown. **c,** Catalytic activity of wild-type SUV420H1 and SUV420H1 truncations on $NCP^{H2A}$ by end-point HMT assays *in vitro*. Each assay was repeated at least three times with similar results. n = 3 independent experiments, two-tailed, unpaired t-test. Adjusted p-values for pairwise ANCOVA comparison of wild type SUV420H1 and each truncation are reported: **** $p < 0.0001$; ns, not significant.

**Table S1**

| Cryo-EM data collection, refinement and validation statistics | | |
|---|---|---|
| | SUV420H1-NCP$^{H2A}$<br>EMDB-36265<br>PDB 8JHG | SUV420H1-NCP$^{H2A.Z}$<br>EMDB-36264<br>PDB 8JHF |
| **Data collection and processing** | | |
| Maginification | 130,000 | 130,000 |
| Voltage (kV) | 300 | 300 |
| Electron exposure (e-/Å$^2$) | 50 | 50 |
| Defocus range (μm) | 1.0-2.0 | 1.0-2.0 |
| Pixel size (Å) | 0.92 | 0.92 |
| Symmetry imposed | C1 | C1 |
| Initial particle image (no.) | 873,129 | 984,971 |
| Final particle image (no.) | 35,448 | 45,377 |
| Map resolution (Å) | 3.58 | 3.68 |
| FSC threshold | 0.143 | 0.143 |
| **Refinement** | | |
| Map sharpening *B* factor (Å$^2$) | -100 | -100 |
| **Model composition** | | |
| Protein residues | 1040 | 1030 |
| Ligands (SAM and $Zn^{2+}$) | 1 SAM | 1 SAM |
| **R.m.s. deviations** | | |
| Bond lengths (Å$^2$) | 0.006 | 0.003 |
| Bong angles (°) | 0.693 | 0.588 |
| **Validation** | | |
| MolProbity score | 1.93 | 2.02 |
| Clash score | 8.48 | 11.25 |
| Poor rotamers (%) | 0.49 | 0.40 |
| **Ramachandran plot** | | |
| Favored (%) | 91.06 | 90.08 |
| Allowed (%) | 8.35 | 9.52 |
| Disallowed (%) | 0.59 | 0.40 |
